# Calcium spikes accompany the cleavage furrow ingression and cell separation during fission yeast cytokinesis

**DOI:** 10.1101/2020.04.07.030379

**Authors:** Abhishek Poddar, Oumou Sidibe, Aniruddha Ray, Qian Chen

## Abstract

The role of calcium signaling during cytokinesis has long remained ambiguous. The studies of embryonic cell division discovered that calcium concentration increases transiently at the division plane just before the cleavage furrow ingression, leading to the hypothesis that these calcium transients trigger the contractile ring constriction. However, such calcium transients have only been found in animal embryos and their function remains controversial. Here we explored cytokinetic calcium transients in the model organism fission yeast. We adopted GCaMP, a genetically encoded calcium indicator, to determine the intracellular calcium level. We validated GCaMP as a highly sensitive calcium indicator which allowed us to capture the calcium transients stimulated by osmotic shocks. To identify calcium transients during cytokinesis, we first identified a correlation between the intracellular calcium level and cell division. Next, we discovered calcium spikes at the start of the cleavage furrow ingression and the end of the cell separation using time-lapse microscopy to. Inhibition of these calcium spikes slowed down the furrow ingression and led to frequent lysis of the daughter cells. We conclude that like the larger animal embryos fission yeast triggers cytokinetic calcium transients which promote the ring constriction and daughter cell integrity (194).

**Highlight summary for TOC:** Calcium rises transiently at the division plane during embryonic cell cytokinesis, but the conservation and function of such calcium transients remain unclear. We identified similar calcium spikes during fission yeast cytokinesis and demonstrated that these spikes promote the contractile ring constriction and the daughter cell integrity (257).

## Introduction

Calcium is an essential secondary messenger in many cellular processes but its role during cytokinesis, the last stage of cell division, remains ambiguous. Eukaryotic cells maintain their intracellular calcium at much lower concentration than that of their extracellular environment. A transient increase of the free calcium level, through either release from the intracellular storage or influx through the plasma membrane, triggers a number of essential calcium signaling pathways (for a review see (Clapham, 2007)).

Although the importance of calcium to cytokinesis had long been known (Arnold, 1975), the calcium transients accompanying cytokinesis were discovered much later and remain poorly understood. Fluck et al. first observed two localized calcium waves at the cell division plane of medaka fish embryos during cytokinesis (Fluck et al., 1991). The first wave initiates just before the cleavage furrow ingression while the second wave appears after the cell separation.

Followed-up studies discovered similar localized increase of calcium in other animal embryos, including those of zebra fish *Danio rerio* and African frog *Xenopus laevis* (Chang and Meng, 1995; Miller et al., 1993; Noguchi and Mabuchi, 2002). However, it remains unexplored whether such calcium transients are universal among all eukaryotic cells. Neither is there a consensus on the exact molecular function of these transients. The prevailing hypothesis (Chang and Meng, 1995; Fluck et al., 1991) holds that these transients can activate myosin light chain kinase (MLCK) to promote the activity of myosin II motor in the contractile ring and trigger the ring constriction (Craig et al., 1983; Scholey et al., 1980). This theory draws a parallel between the role of calcium in cytokinesis to that in smooth muscle constriction (Nishimura et al., 1990). This proposal, although attractive, has remained unproven even in the animal embryos. Adding to the ambiguity, inhibition of the cytokinetic calcium transients in *X. laevis* embryos has produced conflicting results (Miller et al., 1993; Noguchi and Mabuchi, 2002; Snow and Nuccitelli, 1993). To date, both the nature and molecular function of the cytokinetic calcium transients remain undetermined.

Fission yeast *Schizosacchromyces pombe* has emerged as an excellent model organism for the study of cytokinesis in the last twenty years. The molecular mechanism of cytokinesis in this unicellular organism has been largely conserved in higher eukaryotes (for review see (Pollard and Wu, 2010)). Like animal cells, fission yeast assembles an actomyosin contractile ring at the cell division plane (Wu et al., 2003), a process mediated by essential cytokinetic proteins such as Myo2p, Cdc12p and Adf1p (Balasubramanian et al., 1998; Chang et al., 1997; Chen and Pollard, 2011). The assembly of contractile ring is further promoted by a continuous polymerization of actin filaments in the ring (Courtemanche et al., 2016). As in higher eukaryotes, the actomyosin ring constricts to drive the cleavage furrow ingression, assisted by a growing septa (Laplante et al., 2015; Mishra et al., 2013). The terminal step of cell separation (for review see (Garcia Cortes et al., 2016; Sipiczki, 2007) requires degradation of the septa (Cortes et al., 2002; Cortes et al., 2012; Liu et al., 1999; Munoz et al., 2013) as well as proper regulation of the turgor pressure (Atilgan et al., 2015; Proctor et al., 2012).

Although it still remains a challenge to quantify the free calcium concentration in a live cell, the rapid development of genetically encoded calcium indicators (GECI) over the last twenty years has made it far easier than before. Compared to the more widely used synthetic probes, GECIs can be expressed in vivo without intrusive probe-loading procedure. This is particularly advantageous in such genetically modifiable systems as fission yeast. Although most GECIs still trail the chemical probes in their sensitivity, some of them such as GCaMP (Nagai et al., 2001), which we chose for this study, have come close. This single fluorophore sensor consists of three domains: a circularly permuted enhanced green-fluorescent protein (cp-EGFP), a calcium-binding calmodulin motif and a M13 peptide of human myosin-light chain kinase (MLCK). Binding of calcium to the calmodulin motif dissociates the M13 peptide from cp-EGFP, increases the fluorescence of GCaMP significantly (Nakai et al., 2001). Further modifications of GCaMP have substantially reduced its response time and enhanced its signal-to-noise ratio (Chen et al., 2013; Tian et al., 2009). We picked one of the most sensitive variants, GCaMP6s (Chen et al., 2013), to explore the potential calcium transients during fission yeast cytokinesis.

Few studies have examined the calcium level of fission yeast at the single cell level, even though its calcium signaling pathways have been characterized extensively. Two central calcium-signaling molecules, calmodulin and calcineurin, have both been identified (Moser et al., 1997; Takeda and Yamamoto, 1987; Yoshida et al., 1994). As in animal cells, they play a critical role in a plethora of cellular processes including cytokinesis in fission yeast. In particular, the essential calmodulin Cam1p is required for cell division, underlying the importance of calcium-signaling in this model organism (Moser et al., 1997). Past studies have mostly employed bulk assays to determine the average calcium level of many cells, using either chemical or indirect probes (Deng et al., 2006; Ma et al., 2011). As a result, the regulation of calcium at the sub-cellular level during many processes including cytokinesis has been left undetermined.

In this study we first explored the feasibility of GCaMP-based calcium imaging in fission yeast. We then deployed this probe to capture the calcium transients triggered by various stimuli to validate its effectiveness. Finally, we identified the cytokinetic calcium spikes and examined their importance to cytokinesis.

## Results

### Expression and localization of the genetically calcium indicator GCaMP in fission yeast

To probe the intracellular calcium level of fission yeast cells with fluorescence microscopy, we expressed a single copy of the GCaMP coding sequence at an endogenous locus. A strong constitutive ADH promoter drove the expression of GCaMP6s (referred to as GCaMP) (Fig. 1A). To measure the expression and sub-cellular localization of this calcium indicator, we tagged its C-terminus with another fluorescence protein mCherry. GCaMP-mCherry localized uniformly throughout the cytoplasm and the nucleus but it was excluded from the vacuoles (Fig. 1B). The reporter nevertheless was slightly enriched in the nucleus, compared to the cytoplasm (ratio = 1.5 ± 0.1, average ± S.D., n = 66). The ratio of GCaMP to mCherry intracellular fluorescence varied very little among the cells (Fig. 1C), indicating that the expression of this reporter is homogenous. We concluded that GCaMP can be expressed homogenously in the fission yeast cells and this calcium indicator localizes throughout the intracellular space.

**Figure 1.**
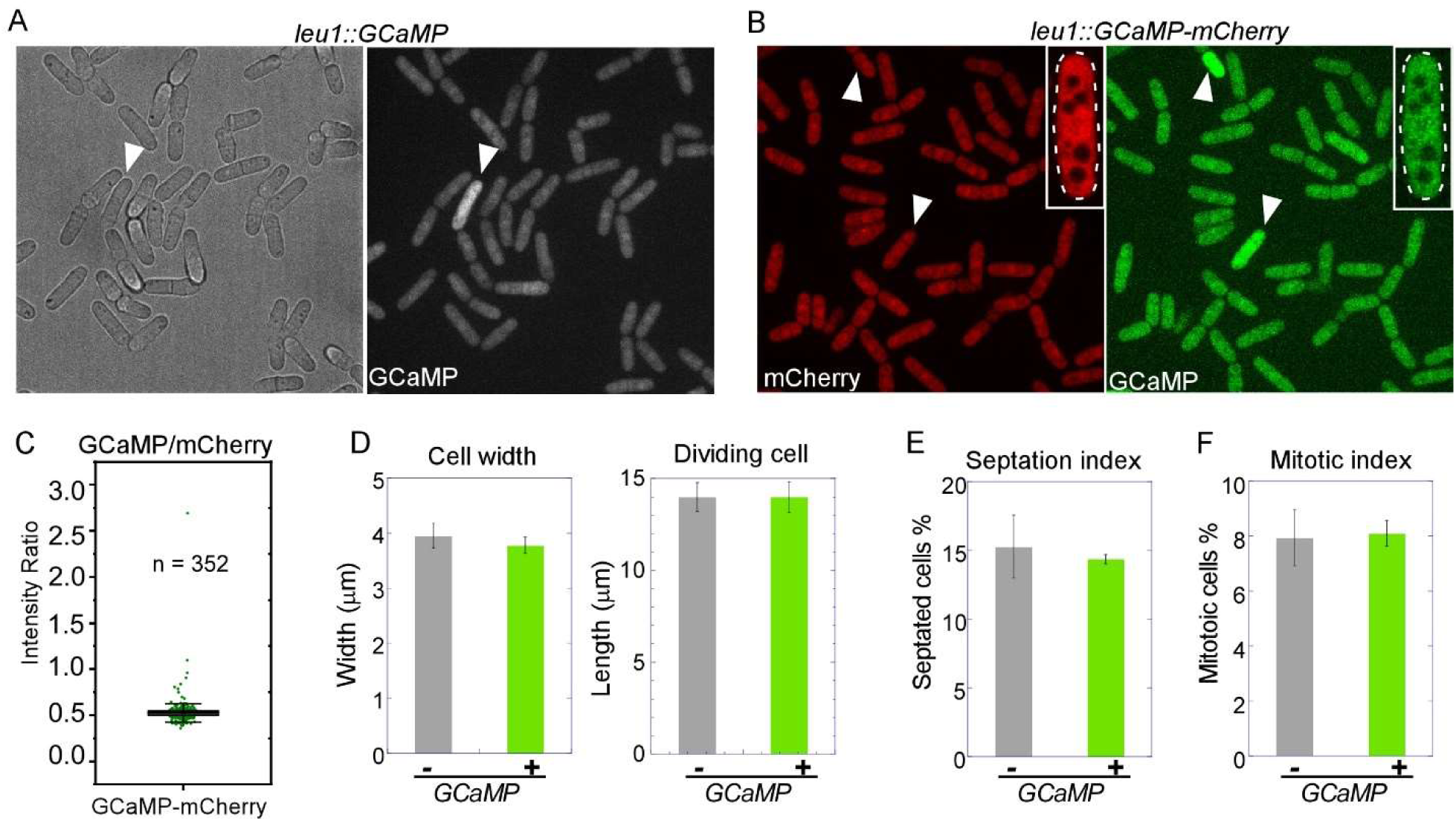
Expression and localization of GCaMP in fission yeast. **(A)** Bright-field (left) and fluorescence (right) micrographs of the cells expressing GCaMP. The intracellular fluorescence was constant for most cells, except for one outlier (white arrowhead). **(B-C)** Localization and expression of GCaMP-mCherry. **(B)** Fluorescence micrographs of the cells expressing GCaMP-mCherry. The insert shows the center slice of the Z-series of a representative cell (magnified, outlined in dashed lines). The intracellular fluorescence of mCherry remained low even in two cells with high GCaMP fluorescence (white arrow heads). GCaMP localized throughout the cytoplasm and nucleus, but it was excluded from the vacuoles. Similar results were found in three biological repeats. **(C)** A box plot showing the ratio of GCaMP:mCherry fluorescence. The expression is homogenous among the cells, resulting in a near uniform ratio with a few outliners with likely high calcium level. **(D-F)** Expression of GCaMP did not produce discernable artifacts. Bars graphs compare the GCaMP-expressing cells to the wild type. **(D)** The width of all cells (left, n > 50) and length (right, n > 50) of dividing cells. No significant differences were found (P > 0.1). **(E-F)** (D) Mitotic index (n > 1000) and (E) septation index (n > 700). No significant differences were found (P > 0.1). The data was from two biological repeats.

We next determined whether the expression of GCaMP perturbs any cellular functions, a potential concern for endogenously expressed reporters. Overall the GCaMP expressing cells exhibited no apparent morphological defects. Their length and width were similar to the wild-type cells (Fig. 1D). Their mitotic and septation indexes were also normal, similar to the wild type (Fig. 1E-F). Moreover, these GCaMP expressing cells did not exhibit any hyper-sensitivity to either calcium, or sorbitol, or EGTA, all of which likely perturb the intracellular calcium homeostasis (Fig. S1). Lastly, cytokinesis in these GCaMP-expressing cells were unperturbed, as determined by time-lapse fluorescence microscopy. In the GCaMP-expressing cells, the contractile ring assembly and maturation took 44 ± 5 mins (average ± S.D., n = 31), starting from the appearance of precursor nodes, similar to the wild-type cells (43 ± 4 mins, n = 40). This was followed by the ring constriction that took 30 ± 3 mins (n = 43), same as the wild type (30 ± 3 mins, n = 70). The last stage of cytokinesis the cell separation required 26 ± 4 mins (n = 29) in the GCaMP-expressing cells, not significantly different from the wild type (25 ± 3 mins, n = 67). These quantitative measurements indicated that expression of GCaMP dose not interfere with cytokinesis either. Since GCaMP does not produce any discernable artifacts, we combined our GCaMP-expressing strain with quantitative microscopy to examine the intracellular calcium level of fission yeast cells.

### Calcium transients in fission yeast cells

To validate GCaMP as an effective calcium indicator, we first determined whether an increase of extracellular calcium alters the fluorescence of GCaMP-expressing cells. In the rich YE5s media, addition of 30mM CaCl_2_ increased average fluorescence of the GCaMP expressing cells by about three folds, compared to those in YE5s (Fig. 2A-B). The increased extracellular calcium concentration also resulted in more frequent fluctuations of the intracellular GCaMP fluorescence (Fig. 2C-D and Supplemental Movie S1). Thus, GCaMP-based imaging is fully capable of detecting alterations of calcium concentration.

**Figure 2.**
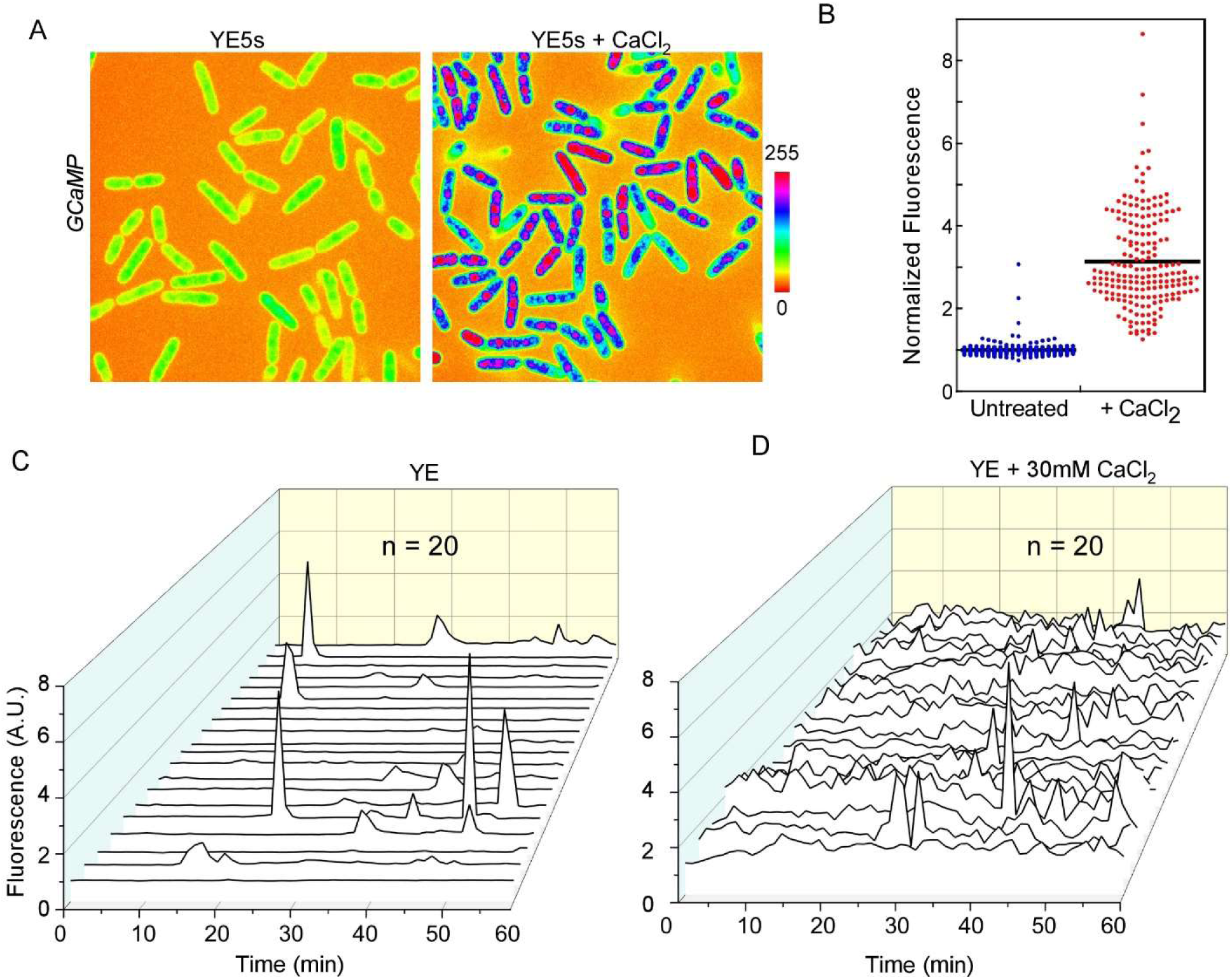
GCaMP responds to the calcium level increase. **(A-B)** The intracellular GCaMP fluorescence increased with added calcium in the media. **(A)** Fluorescence micrographs (spectrum colored) of the GCaMP-expressing cells in either YE5s (left) or YE5s supplemented with 30mM CaCl_2_ (right). Bar represents the intensity scale. **(B)** Dot plot of fluorescence intensities of the cells (n > 190). Black line represents the average. Added calcium increased the average intracellular fluorescence by ∼3 folds. **(C-D)** Calcium homeostasis of fission yeast cells. 3D-line plots show the time courses of intracellular GCaMP fluorescence in either YE5s (C) or YE5s plus 30mM CaCl_2_ (D). The time-lapse movies were recorded at a frequency of 1 frame/min. Increased calcium in the media resulted in more frequent fluctuations of the intracellular calcium level. The data was pooled from two biological repeats.

Next, we employed the GCaMP-based time-lapse imaging to capture rapid changes of intracellular calcium level, the calcium transients. The resting fission yeast cells produced only sporadic calcium transients (Fig. 2C). We applied various stimuli through microfluidics to produce more transients. First, we used hypo-osmotic shock, one of the best-known stimuli of calcium transients in yeast (Batiza et al., 1996; Denis and Cyert, 2002). We found highly synchronized calcium transients, captured by an increase of GCaMP fluorescence, in the cells under such shock (Fig. 3A and Supplemental Movie S2). The intracellular calcium rose quickly following the shock and peaked 2 mins after the osmotic shock on average. These calcium transients accompanied the expansion of cell volume closely (Fig. 3B-C). Average amplitudes of the calcium increase were proportionally to the strength of hypo-osmotic shocks (Fig. 3D). Secondly, we applied hyper-osmotic shocks to the cells. This surprisingly did not elicit detectable calcium transients (Fig. 3E and Fig. S2A-B). Lastly, we abrupted increased the extracellular calcium concentration to stimulate the cells. This also provoked strong calcium transients, but they (QC) appeared only ∼5 mins after the infusion of calcium (Fig. 3F). As a result of the stimulation, the average intracellular calcium concentration remained elevated after more than 20 mins. This response to the external calcium shock confirmed a similar observation made through the bulk assays (Deng et al., 2006). Our results demonstrated that the GCaMP-based time-lapse imaging can capture the calcium transients in fission yeast cells, making it entirely feasible to identify the cytokinetic calcium transients.

**Figure 3.**
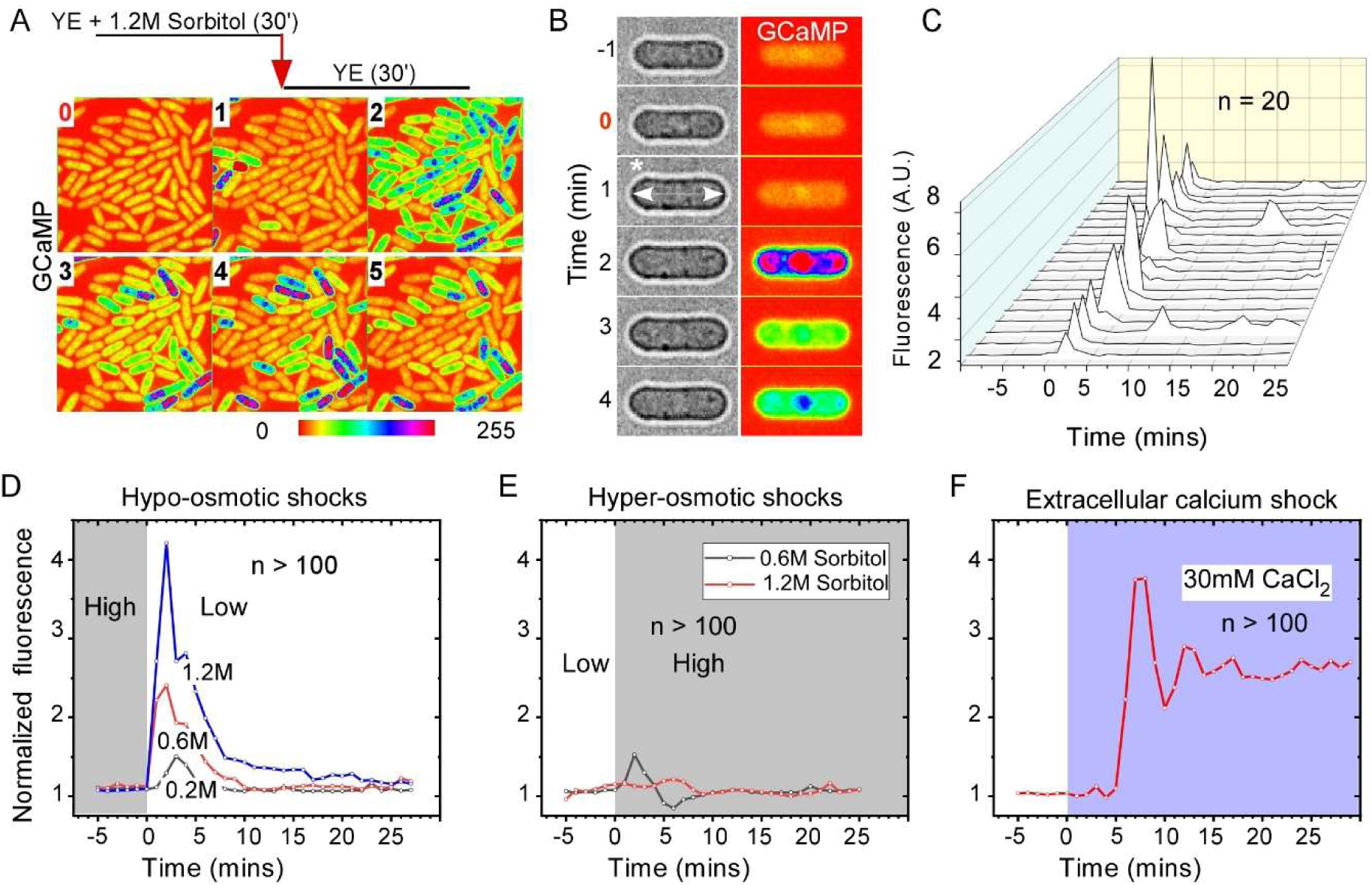
Osmotic shocks trigger calcium transients. **(A-C)** Hypo-osmotic shock triggered calcium transients in fission yeast cells. The GCaMP expressing cells, trapped in a microfluidic chamber, were first equilibrated in YE5s supplemented with 1.2M sorbitol for 30 mins before the infusion of YE5s (time zero). **(A)** Time-lapse micrographs (spectrum-colored) of a representative field in the chamber. The hypo-osmotic shock triggered a synchronized calcium spike in the cells. **(B)** Time-lapse micrographs of a representative cell. In the media with 1.2M sorbitol, the cells appeared shrunk. After the shock (time zero), the cell quickly expanded (asterisk), followed by calcium transient (right, spectrum-colored). Number: time in minutes. **(C)** 3D-line plots of the time courses of GCaMP fluorescence in twenty representative cells. The shock triggered synchronized calcium transients in every cell. Representative results from two biological repeats are shown. **(D-F)** Average time courses of the intracellular GCaMP fluorescence during either hypo-osmotic (D), or hyper-osmotic (E), or extracellular calcium (F) shocks applied through microfluidics. These stimuli triggered distinct change of intracellular calcium. **(D)** The cells were first equilibrated in YE5s with various concentrations of sorbitol for 30 mins before the infusion of YE5s media (time zero). The average amplitudes of calcium level increase were proportional to the strength of the shocks. **(E)** The cells were first equilibrated in YE5s media for 30 mins before the infusion of YE5s media supplemented with various concentrations of sorbitol (time zero). No significant change of calcium was triggered by hyper-osmotic shock. **(F)** The cells were equilibrated in YE5s media for 30 mins before the infusion of YE5s plus 30mM CaCl_2_ (time zero). The addition of extracellular calcium triggered a delayed increase of intracellular calcium. All data is pooled from two biological repeats.

### Identification of calcium spikes during cytokinesis

As the first step to determine the link between calcium and cytokinesis, we analyzed the intracellular calcium level in a large number of asynchronized cells. Dividing fission yeast cells can be identified through their increased length. These rod-shaped cells grow by tip extension, starting to grow from a length of ∼7 µm after birth. They expand throughout interphase until stopping at a length of ∼14 µm at the mitotic entry (Mitchison and Nurse, 1985). We found that the calcium level was constant among most cells regardless of their length (n = 407) (Fig. 4A). However, the calcium centration of a few cells was significantly higher than the average (>110%). The length distribution of these “outliers” was interestingly bimodal. Two peak fractions are ∼14 µm and ∼8 µm respectively (Fig. 4B), roughly equaling to the length of the dividing and new-born cells respectively. The existence of the outlier cells with high calcium level provided the first hint of calcium transients during cell division.

**Figure 4.**
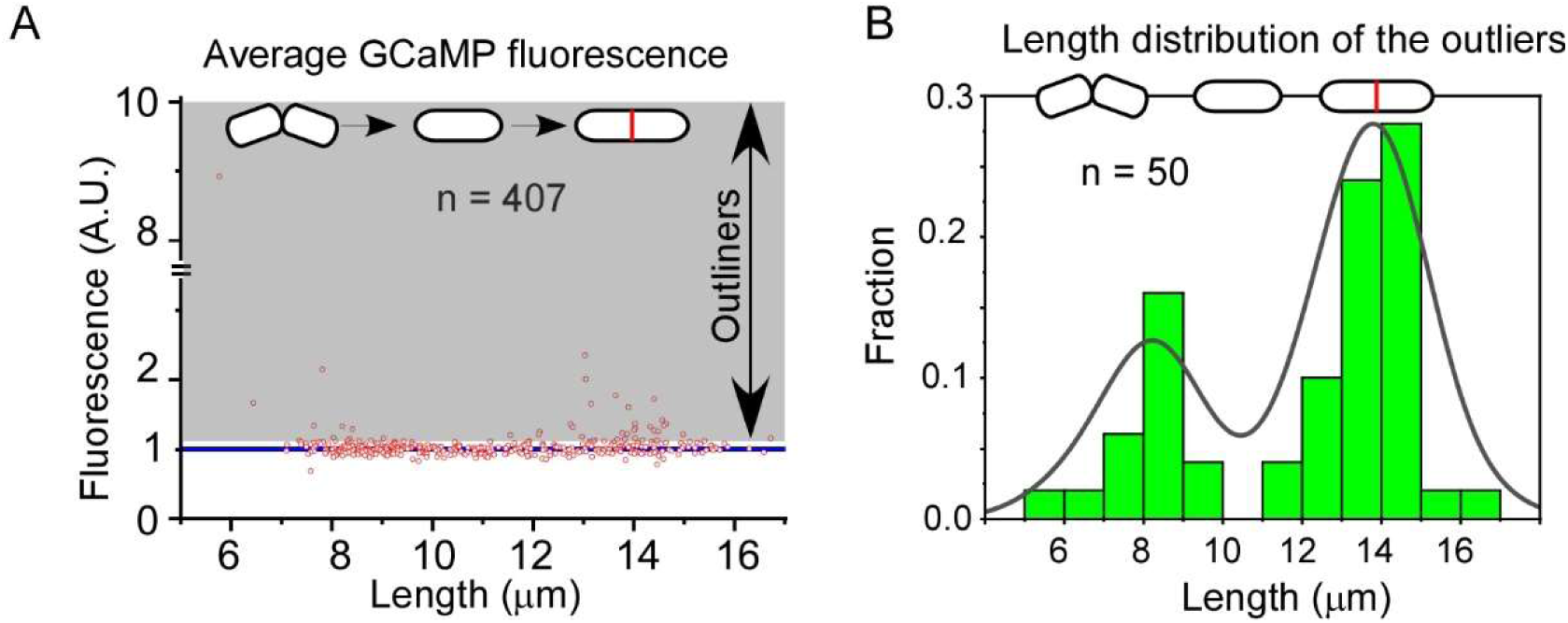
Correlation between intracellular calcium level and cell division. The intracellular calcium level in asynchronized cells, based on the GCaMP fluorescence. Top: cartoon representations of fission yeast cells at birth, growth, and division. **(A)** A scatter plot showing the relationship between the intracellular calcium level and the cell length. The calcium level of most cells was uniform (blue line). Only a few outlier cells (50/407) exhibited elevated (shaded area, >110% of the average) calcium level. **(B)** A histogram showing the distribution of the cell length among the outliers. The black line represents the best fit of a bimodal distribution. Two peak fractions represent the cells of 8-9 µm, and those of 14-15 µm, roughly corresponding to newly-born and dividing cells respectively. The data is pooled from four biological repeats.

Next, we examined the temporal regulation of intracellular calcium in the dividing cells through time-lapse microscopy. To facilitate our analysis, these cells expressed both GCaMP and well-established fluorescence protein markers of cytokinesis. First, we looked closely at the calcium level during mitosis, by imaging the cells expressing both GCaMP and Sad1p-GFP, a marker of spindle pole bodies (SPB). In fission yeast, the separation of SPBs is concomitant with the start of prometaphase and anaphase B ends ∼30 mins after (Fig. S3A) (Wu et al., 2003). During this period, we captured very few calcium transients. About 34 mins after the separation of SPBs, the calcium level started to rise quickly throughout the intracellular space, which we termed calcium spikes. These spikes appeared to be well-synchronized, coinciding with telophase as well as early cytokinesis (Fig. S3B-C). Next, we determined whether these calcium spikes initiated exactly at the beginning of furrow ingression as those calcium transients did in the fish embryos (Chang and Meng, 1995; Fluck et al., 1991). We imaged the cells expressed GCaMP and a contractile ring marker Rlc1p-tdTomato, the regulatory myosin light chain (Fig. 5A). Our movies recorded very few calcium spikes during the assembly and maturation of the contractile ring (Fig. 5A-B). In comparison, our time-lapse microscopy consistently captured calcium spikes accompanying the start of ring constriction in most dividing cells. The spikes also appeared frequently during the constriction. We called them “constricting spikes” (Fig. 5B-C and Supplemental Movie S3). In these cytokinetic cells, the average intracellular calcium level increased ∼1.7 fold within 1 ± 3 min (average ± S.D., n = 64) of the start of the cleavage furrow ingression (Fig. 5D).

**Figure 5.**
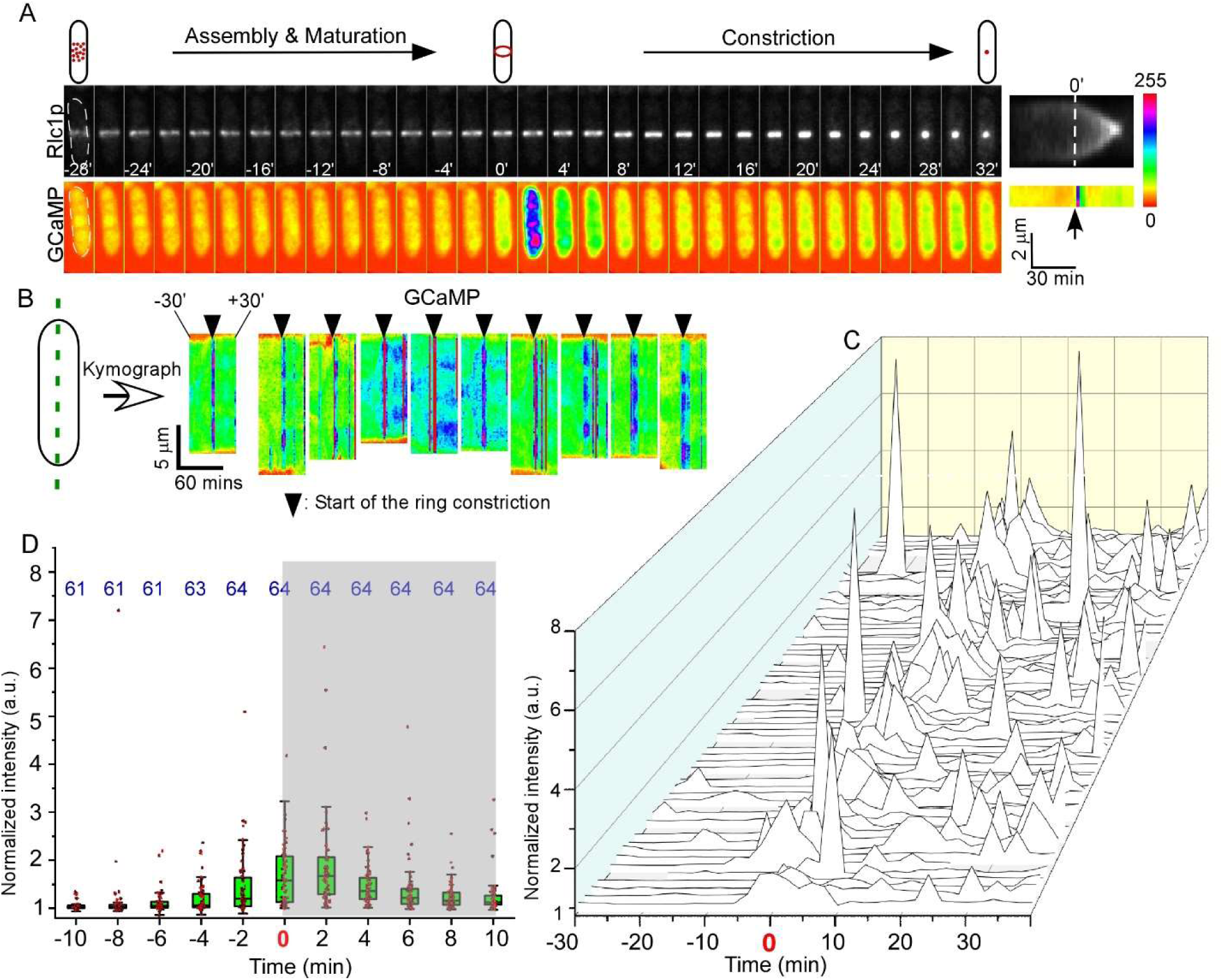
Calcium spikes accompany the cleavage furrow ingression. **(A-B)** The calcium spikes accompanied the contractile ring constriction. **(A)** Left: Time-lapse micrographs of a dividing cell expressing both Rlc1p-tdTomato (gray, top) and GCaMP (spectrum-colored, bottom). Number: time in minutes after the start of ring constriction. Interval = 2 mins. Right: Kymograph of the contractile ring (top) and the GCaMP fluorescence, with the dash line marking the start of ring constriction. A calcium spike initiated at time zero and peaked at +2 min. Bar represents the relative scale of intracellular calcium level. **(B)** Kymographs of GCaMP fluorescence in ten representative cells from −30 mins to +30 mins, relative to the initiation of ring constriction. **(C-D)** Quantitative analysis of the intracellular calcium level during the ring constriction. Data is pooled from three biological repeats. **(C)** 3-D line plots showing the time courses of calcium level in representative cells (n = 64). **(D)** Box plots of intracellular calcium level in the cells shown in C. The number on top indicates the number of observations made at each time point. The start of ring constriction was accompanied by emergence of calcium spikes. The average intracellular calcium level started to increase significantly at −4 mins and peaked at +2 mins, relative to the start of ring constriction.

Besides the dividing cells, the intracellular calcium of new-born cells appeared to be high as well, prompting us to examine the calcium spikes during cell separation. Although few calcium spikes were captured following the ring closure and before the cell separation, spikes were found in most cells following the cell separation (Fig. 6A). These separating spikes were concomitant with the initial appearance of new ends. They distributed asymmetrically between the two daughter cells (Fig. 6B-C). Among the cells (n = 73) in which at least one separating spike was captured, 56% triggered the spike in only one daughter cells. The other 44% triggered the calcium spikes that oscillated between the two daughter cells (Fig. 6B-C). On average, the separating spikes increased the intracellular GCaMP fluorescence by ∼1.7 folds (Fig. 6D-E). The calcium level peaked ∼2 mins after the end of cell separation (Supplemental Movie S4). We observed similar constricting and separating spikes in the dividing cells inoculated in the synthetic EMM5s media (Fig. S3D-E), further confirming the existence of these cytokinetic spikes.

**Figure 6.**
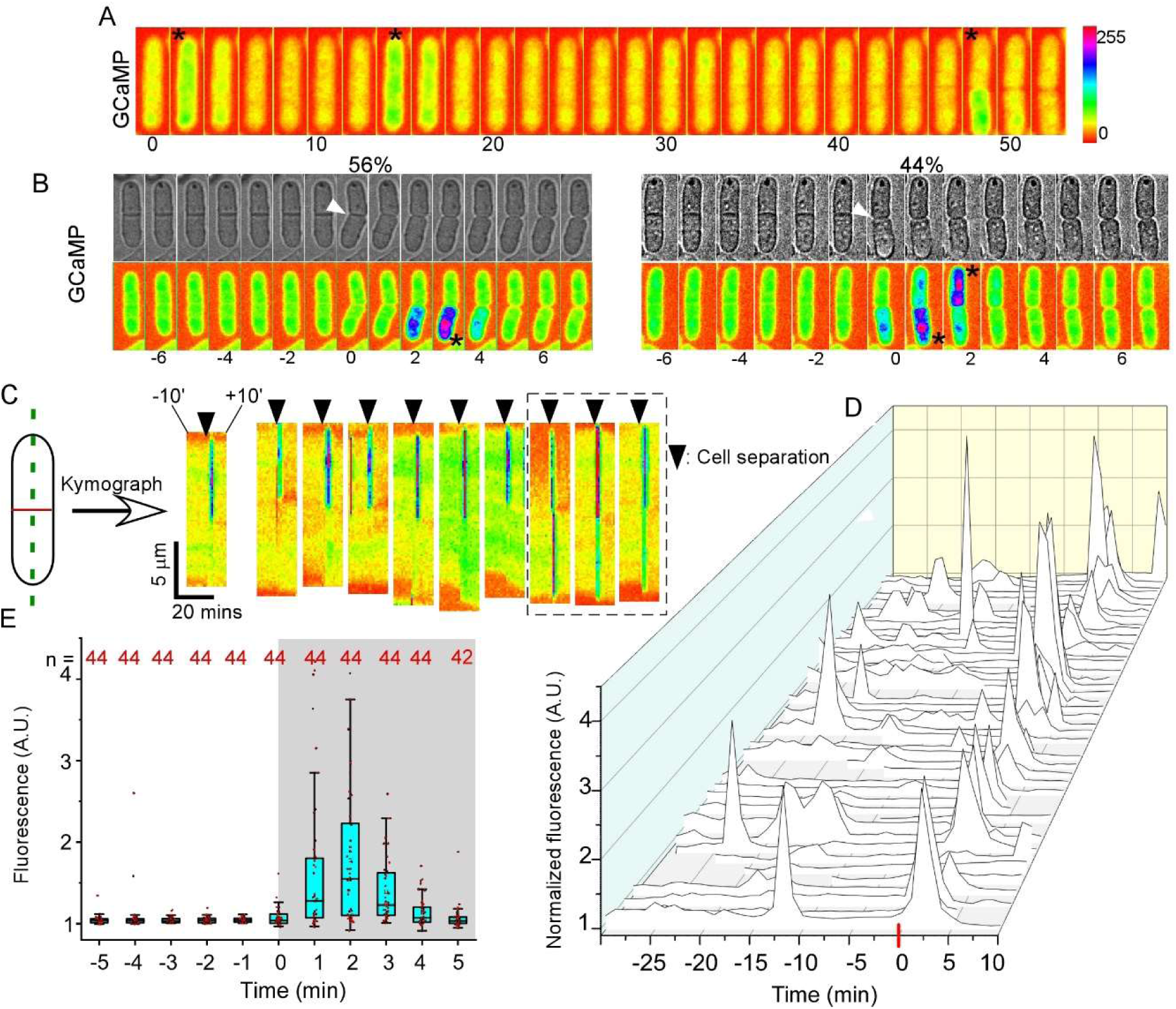
Calcium spikes accompany the cell separation. **(A-C)** Cytokinetic calcium spikes during cell separation (white arrowhead). Asterisk: the peak of a spike. Bar: the relative scale of intracellular calcium level. **(A)** Time-lapse micrographs of a cell expressing both GCaMP (spectrum-colored) and Rlc1p-tdTomato (not shown). Three calcium spikes (asterisk) are detected in this cell, including one following the start of ring constriction (time zero), the second one during the ring constriction and the third one after the cell separation (arrowhead). Number: time in minutes. Interval = 2 mins. **(B)** Time-lapse micrographs of two cells expressing GCaMP (bottom). Interval = 1 min. Number: time in minutes after the cell separation (time zero, arrowhead). The spikes appeared either in just one daughter cell (top) or oscillate between two daughter cells (bottom). Arrowhead: completion of the cell separation. (**C)** Kymograph of separating spikes in ten representative cells. Dashed box: the separating spikes oscillating between two daughter cells. **(D-E)** Quantification of the intracellular calcium level during the separation. The time-lapse movie was captured at a frequency of 1 frame/min. The data is pooled from four biological repeats. **(D)** 3-D line plots showing the time courses of intracellular calcium level in the dividing cells (n = 44). A highly synchronized calcium spike followed the cell separation (time zero) in most cells. **(E)** Box plots of the calcium level of the cells shown in D. Number of observations at each time point is indicated on the top. The average calcium level started to increase at 0 min and peaked at +2 min.

To better understand the spatiotemporal regulation of these cytokinetic calcium spikes, we characterized them by fast imaging. We increased the frequency of time-lapse acquisition by 40 folds, from one frame every two minutes to one frame every three seconds. We only captured a limited number of constricting and separating spikes in this way (Fig. 7A and D). Close examination of the spikes found that they increased intracellular calcium level most significantly at the tip, the peri-nuclear space, and the division plane (Fig. 7B and E). All of them exhibited a lifespan of 50-200 seconds (Fig. 7C and F). Their amplitudes were highly variable (Fig. 7C and F). Some constricting spikes exhibited a more than 10-fold increase of GCaMP fluorescence above the baseline level (Fig. 7C). We concluded that the cytokinetic calcium spikes rise throughout the cytoplasm and they are highly heterogeneous.

**Figure 7.**
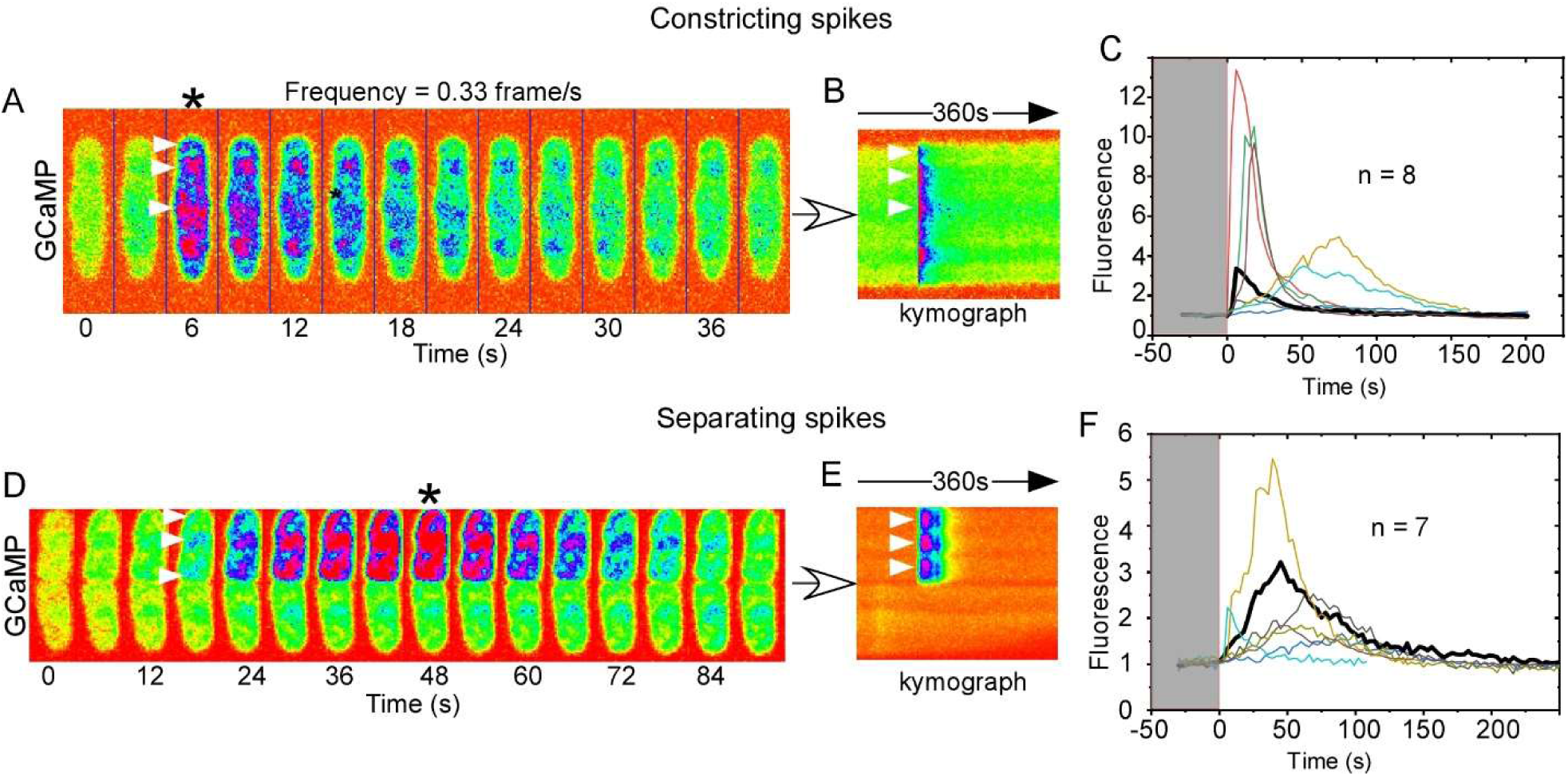
Spatiotemporal regulation of cytokinetic calcium spikes. The spikes were captured at a frequency of one frame every three seconds during the time-lapse microscopy. Representative constricting (A-C) or separating spikes (D-F) are shown. **(A, D)** Time-lapse micrographs (spectrum-colored) of two GCaMP-expressing cells during cytokinesis. Arrowhead: hots spot of calcium increase in the cytoplasm. Number: time in seconds after the initial rise of calcium (Time zero). Asterisk: the peak of calcium level. **(B, C)** Fluorescence kymograph of the spikes. Calcium increased most significantly at the cell tips, the peri-nuclear area and the cell division plane during spikes. **(C, F)** Time course of cytokinetic spikes including the one shown on the left (thick black line). The cytokinetic calcium spikes are highly heterogeneous with various amplitudes and lifespans. Data is pooled from three biological repeats.

### Functions of the fission yeast cytokinetic calcium spikes

The close temporal correlation between the calcium spikes and cytokinesis prompted us to investigate whether these spikes are required for cytokinesis. To this end, we depleted the extracellular calcium from the media, hypothesizing that the calcium influx contributes to the spikes. The calcium depletion was only temporary to minimize the effects on the other calcium-dependent processes. The cells were observed in either EMM5s media without any supplemented calcium or YE5s media supplemented with low concentrations of EGTA. At such concentrations, we found that EGTA did not inhibit cell growth strongly (Fig. S4A). In the calcium-depleted media, the average intracellular GCaMP fluorescence decreased only slightly, ∼13% in YE5s plus 2 mM EGTA and ∼3% in the calcium-free EMM5s media (Fig. 8A-B). However, the fraction of outlier cells decreased significantly under either condition (Fig. 8A-B). Although the calcium spikes were no longer as visible under either conditions, quantitative image analysis revealed that the total number of cytokinetic spikes was not reduced eve though their amplitudes decreased dramatically (Fig. 8C-D and Fig. S4B-E). We concluded that the cytokinetic calcium spikes can be inhibited by depleting extracellular calcium from the media.

**Figure 8.**
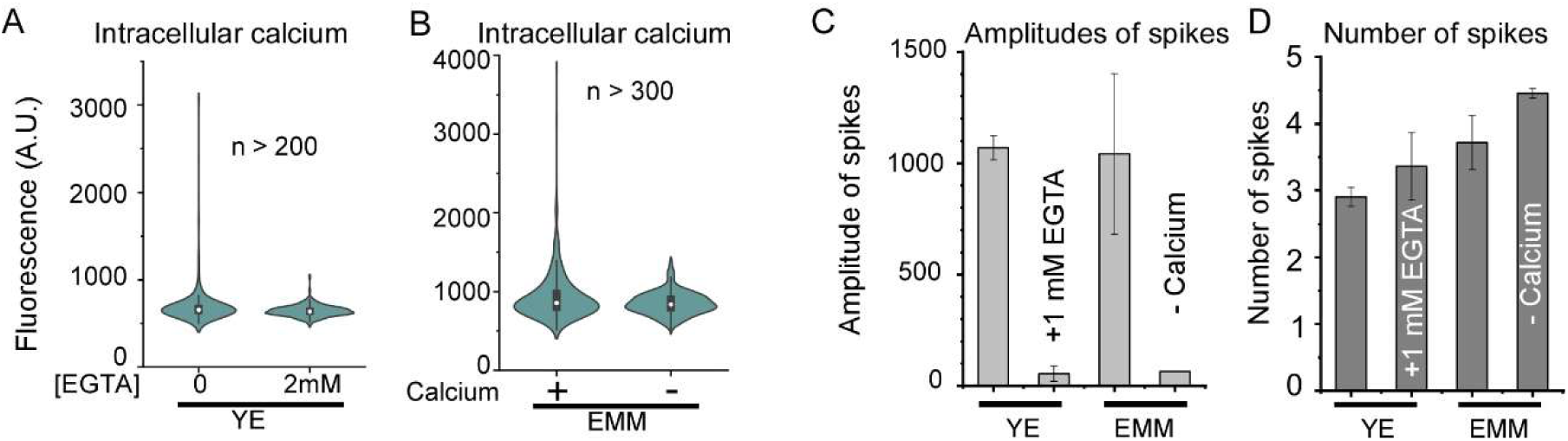
Depletion of extracellular calcium inhibits the cytokinetic calcium spikes. Either supplement of EGTA or removal of calcium from the media inhibited the calcium spikes. **(A-B)** Violin plots showing the intracellular calcium level in the cells. Depletion of calcium from the media reduced the average intracellular calcium level only slightly (13% by EGTA and 3% by the calcium-free EMM media) but it greatly reduced the number of outlier cells with high calcium level. **(C-D)** Bar graphs showing the average amplitude (E) and number (F) of calcium spikes during the ring constriction (n = 20). The amplitude and number of spikes are quantified through thresholding using custom-written algorithm as described in the Materials and Methods. Depletion of extracellular calcium reduced the average amplitude of calcium spikes significantly (P < 0.001), but not the number of peaks. The data is pooled from two biological repeats.

We next examined cytokinesis in the cells whose calcium spikes are inhibited through time-lapse microscopy. We found that the assembly and maturation of the contractile ring took slightly longer time in the calcium-free EMM media, compared to the EMM media (Fig. 9A and C). However, constriction of the contractile ring slowed down much more dramatically (30%, Fig. 9E and Supplemental Movie S5). Similarly, 1-2 mM EGTA did not alter the assembly and maturation of the ring significantly (Fig. 9B and D) but the constriction rate was inhibited by EGTA in a dosage-dependent manner. The rate was down ∼50% in YE5s plus 2mM EGTA, compared to YE5s (Fig. 9E). When EGTA concentration was further increased to 5mM, most contractile ring (90%, n = 30) disintegrated without completing the constriction (Fig. S5A-B). We also examined the cell separation when the calcium spikes are inhibited. Surprisingly, the duration of cell separation remained normal when the extracellular calcium was depleted (Fig. S5C-D). Although no daughter cells lysed in the calcium-free EMM5s media, 2 mM EGTA induced more than one third (35 ± 5 %, n = 103) of the daughter cells to lyse following the separation (Fig. 9G-H). More cells lysed when the concentration of EGTA increased to 5 mM (data not shown). Further depletion of intracellular calcium by deletion of *pmr1* resulted in 54% of daughter cells lysed in the calcium-free EMM5s media even though no mutant cells lysed in the regular EMM5s. Pmr1p is a ER calcium ATPase that replenish the internal storage of calcium (Cortes et al., 2004). We concluded that the cytokinetic calcium spikes also likely promote the integrity of separating cells.

**Figure 9.**
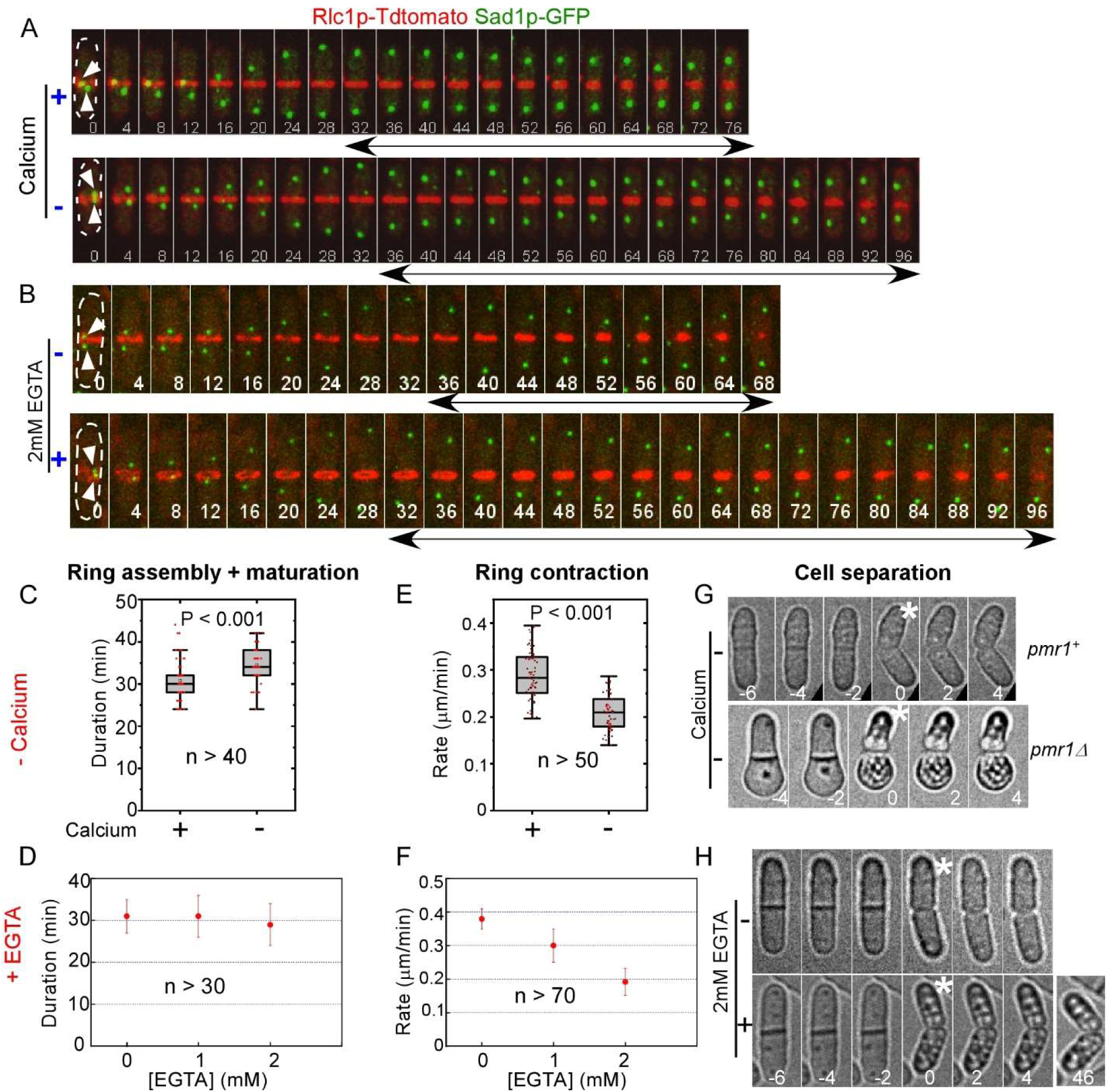
Temporary depletion of extracellular calcium slows down the contractile ring constriction and leads to daughter cell lysis. **(A-F)** Temporary depletion of extracellular calcium reduced the ring constriction rate, but it had little effect in the ring assembly and maturation. **(A-B)** Temporary depletion of extracellular calcium slowed down the ring constriction. Time-lapse micrographs of dividing cells expressed both Sad1-GFP (green) and Rlc1p-tdTomato (red). Number: time in minutes after the separation of SPBs (arrowhead). Dash line marks the outline of cells. **(A)** Cells in either regular EMM5s (top) or the calcium-free EMM5s media (bottom). **(B)** Cells in either YE5s (top) or YE5s plus 2mM EGTA media (bottom). **(C, D)** Duration of the ring assembly plus maturation. **(E, F)** The ring constriction rates. **(G-H)** Temporary depletion of extracellular calcium increased the frequency of cell lysis among the daughter cells. Time-lapse micrographs of separating cells. Number: time in minutes after the cell separation. **(G)** The wild-type (top) or *pmr1Δ* (bottom) cells in the calcium-free EMM5s media (bottom). Although the wild-type cells (n > 100) did not lysis during the separation, 54% of the mutant cells lysed (asterisk) (n = 57). **(H)** The wild-type cells in either YE5s (top) or YE5s plus 2mM EGTA (bottom). About 35% of daughter cells lysed after the separation in the presence of EGTA (n > 100). Data is pooled from two biological repeats.

## Discussion

Overall, we demonstrated that GCaMP-based imaging will allow fission yeast to serve as a powerful model organism for the study of calcium transients during various cellular processes. Using this method, we discovered the cytokinetic calcium spikes in this unicellular model organism for the first time. These calcium transients are similar to those first uncovered in animal embryos. They likely play a critical role in promoting the ring constriction and the daughter cell integrity.

### GCaMP-based calcium imaging of fission yeast

To our knowledge, our study is the first one to examine the feasibility of GCaMP-based calcium imaging in fission yeast cells. We are fortunate that fission yeast is quite tolerant to the expression of this calcium indicator. It does not perturb many cellular processes that we examined even when it is expressed constitutively. More importantly, GCaMP is also highly sensitive as a probe for time-lapse microscopy, allowing us to capture calcium transients at relatively low sampling rate. Compared to synthetic calcium probes, GCaMP exhibited two key advantages in fission yeast. First, unlike the synthetic probes such as Calcium Green (Chang and Meng, 1995), the intracellular concentration of GCaMP can be maintained at a constant level through the homogenous expression. Combined with quantitative microscopy, this largely eliminated the heterogeneity of fluorescence intensities among cells. In contrast, intracellular concentration of the synthetic probes can be highly variable among cells. This advantage was exemplified by the identification of outliers with high calcium among a large number of cells. The other advantage of GCaMP is its more uniform sub-cellular localization, compared to the synthetic probes. This allows detection of calcium transients throughout the cytoplasm as well as the nucleus.

When combined with yeast genetics, GCaMP can be a very versatile tool for studying calcium signaling. As an example, we constructed both GCaMP and GCaMP-mCherry for this study, both of which have the potential to be employed in the future studies. We employed GCaMP not GCaMP-mCherry throughout this study, primarily to preserve the spectrum for imaging the fluorescence protein markers of cytokinesis. Nevertheless, the tandem reporter can be used for ratio-metric imaging of intracellular calcium, potentially providing higher signal to noise ratio. Overall, application of GCaMP shall make fission yeast an excellent model organism to study calcium transients and homeostasis in non-excitable cells.

### Evolutional conservation of the cytokinetic calcium transients

Our study provides fresh evidences for the existence of calcium transients in an unicellular organism. Similar to the earlier studies (Chang and Meng, 1995; Fluck et al., 1991; Noguchi and Mabuchi, 2002), we employed live fluorescence microscopy to determine the temporal correlation between the calcium transients and cytokinesis. With the availability of fluorescence protein markers of cytokinesis, we now determined the temporal regulation of these spikes with more precision. In addition to time-lapse microscopy, we sought out an alternative approach to illustrate this close correlation. We analyzed just snapshots of many GCaMP-expressing cells to reveal the relationship between cell division and intracellular calcium level. Compared to the movies, the snapshots examined all cells regardless of their cell-cycle stage, presenting a less biased view of the potential calcium change in them. Combined with the previous studies of animal embryos, our results strongly support that a temporal increase of calcium during cytokinesis may be evolutionally conserved.

The temporal regulation of fission yeast cytokinetic calcium spikes bears strong similarities to those found in the animal embryos. The constricting spikes of fission yeast are comparable to the first “furrowing wave” observed in the animal embryos (Fluck et al., 1991) which initiates just as the cleavage furrow starts to ingress (Chang and Meng, 1995; Noguchi and Mabuchi, 2002). The separating spikes are comparable to the second “zipping wave” of the embryos, starts just when the two daughter cells separate (Fluck et al., 1991; Noguchi and Mabuchi, 2002). Both the calcium waves in the embryos and the spikes in fission yeast can last for minutes, far longer than other known calcium transients (Jaffe and Creton, 1998). Although no calcium spikes have been identified yet during budding yeast cytokinesis, Carbo et al. observed higher frequency of calcium bursts during G1 to S phase transition in synchronized cells (Carbo et al., 2017), which may be similar to the separating calcium spikes in of fission yeast. Overall, our study suggests that the regulatory mechanism of calcium during cytokinesis may have been conserved as well.

There are also key differences between the calcium spikes and those embryonic calcium waves. Chief among them is the spatial regulation of these calcium transients. In fission yeast, the calcium spikes propagate globally throughout the cytoplasm and nucleus. In contrast, the calcium waves of animal embryos are restricted to the cleavage furrow (Chang and Meng, 1995; Fluck et al., 1991) or its surrounding region (Noguchi and Mabuchi, 2002). This difference may be due to varying sensitivity of the calcium probes. It is more likely that this distinction is due to the relatively small size of fission yeast cells, measuring only 4 µm wide and 14 µm long, compared to the embryos of hundreds of microns. So far, we found no evidence that the constricting spike of fission yeast triggers the contractile ring constriction, even though there are evidences that the furrowing waves trigger the ring constriction (Chang and Meng, 1995; Fluck et al., 1991; Miller et al., 1993). Lastly, the asymmetry and oscillation of the separating spikes are also unique to fission yeast. The oscillating behavior of the spikes is particularly striking. It may be linked to the unbalanced turgor pressure between the two daughter cells during the separation.

### Mechanism of the calcium spikes and their potential roles

Although the regulatory mechanism of these cytokinetic calcium spikes remains to be explored, our data suggests that they likely draw calcium from both the influx and internal release. Inhibition of the influx through depleting extracellular calcium significantly inhibited the spikes but not completely. Under such condition, the cytokinetic calcium spikes still occurred albeit with greatly diminished amplitudes. This points to the ER- or vacuole-stored calcium as the other likely sources for the spikes. Supporting this hypothesis is our observation that EGTA exhibited a stronger inhibitory effect to cytokinesis, compared to the calcium-free EMM medium. EGTA may have chelated the intracellular calcium slowly over long-term incubation. This is consistent with the lysis of *pmr1Δ* mutant cells in the calcium-free medium. In comparison, the embryonic calcium waves draw only from the internal store (Chang and Meng, 1995; Miller et al., 1993). It remains unclear what are the ion channels mediating the cytokinetic calcium transients. Fluck et al. has first proposed the potential role of tension-sensing calcium channels which can be activated during cytokinesis (Fluck et al., 1991). This remains a feasible mechanism considering increased membrane tension on the cleavage furrow. Release of internal calcium could be through the store-operated calcium release (SOCE) channels (Chan et al., 2016; Chan et al., 2015). Nevertheless, none of these cytokinetic ion channels has been identified.

It remains unknown how the calcium spikes promote the contractile ring constriction and the daughter cell integrity. One likely explanation is that a transient increase of intracellular calcium may increase the activity of type II myosin in the ring through the phosphorylation of myosin regulatory light chain (Chang and Meng, 1995; Fluck et al., 1991). Alternatively, the spikes could stimulate the septum biosynthesis through activating the calcium-dependent calcineurin (Cadou et al., 2013; Martin-Garcia et al., 2018; Yoshida et al., 1994). Lastly, the calcium spikes may adjust the turgor pressure which is required for the separation (Abenza et al., 2015; Proctor et al., 2012). Further studies will be required to determine the mechanism of calcium signaling during cytokinesis.

### Experimental procedures

#### Yeast genetics

We followed the standard protocols for yeast cell culture and genetics. Tetrads were dissected with a Spore+ micromanipulator (Singer, UK). Yeast cells were transformed with the lithium-acetate method. The pmr1Δ mutant is a commercially available fission yeast deletion library (Bioneer, South Korea). All the strains used in this study are listed (Table 1).

To construct the GCaMP expressing strain, we amplified the coding sequence from *pCMV-GCaMP6s* (Addgene, Plasmid #40753) through PCR using primers P589 and P590. The DNA fragment was subcloned into a previously described plasmid *pFA6a-KanMX6-Padh1-Tadh1* (Chen and Pollard, 2013). The GCaMP6s ORF, together with the Adh1 promoter and terminator, were integrated into the *leu1* locus through PCR-based homologous recombination using primers P413 and P414 (Bahler et al., 1998). The resulting strain was confirmed through both PCR using primers P589 and P416 and the Sanger sequencing using primer P668. The GCaMP-mCherry strain was constructed through homology-based PCR using primers P647 and P648. The genome locus was sequenced to confirm the insertion of mCherry coding sequence.

#### Fluorescence microscopy

For time-lapse microscopy, the cells were first inoculated in liquid YE5s media at 25°C for two days before they were harvested during the exponential growth phase at a density between 5*10^6^/ml to 1.0*10^7^/ml. 20 µl of the cell culture were spotted onto the glass-coverslip (#1.5) in a 10 mm petri dish (Cellvis, USA). The coverslip was pre-coated with 50µl of 50-100 µg/ml of lectin (Sigma, L2380) and allowed to dry for 3-4 hours in a 30°C incubator. After 10 min at room temperature, most cells attached to the coverslip in the petri dish. 2ml of YE5s media was then added to the dish before microscopy. When chelation of calcium in media is required, 2ml of YE5s supplemented with EGTA was added instead.

For the experiments in the calcium-free media, we prepared the synthetic EMM media by omitting CaCl_2_ (100µM) and replacing calcium pantothenate (Vitamin B5) with sodium pantothenate (Cayman Chemical, Cat# NC1435671). The cells were first inoculated in YE5s media for 1 day, followed by another ∼15 hours in EMM5s. After harvesting the exponentially growing cells, we washed them three times with the calcium-free media before proceeding to microscopy.

We employed a spinning disk confocal microscope for the time-lapse microscopy. The microscope is an Olympus IX71 unit equipped with a CSU-X1 spinning disk unit (Yokogawa, Japan). The motorized stage (ASI, USA) includes a Piezo Z Top plate for acquiring Z-series. The images were captured on an EMCCD camera (IXON-897, Andor) controlled by iQ3.0 (Andor). Solid-state lasers of 488 nm and 561 nm were used in the fluorescence microscopy, at power of no more than 2.5mW. Unless specified, we used a 60x objective lens (Olympus, Plan Apochromat, NA = 1.40). Typically, a Z-series of 8 slices at an interval of 1 µm was captured at each time point during the time-lapse microscopy. The temperature was maintained at ∼22 ± 2°C in a designated room for microscopy. To further minimize environmental variations, we typically imaged both control and experimental groups with randomized order in the same day.

For visualization of septum, we fixed the cells with 4% formaldehyde (EMS, USA) before staining them with 1 µg/ml of calcofluor white (Sigma, Cat# 18909). For visualization of nuclei, we fixed the cells with 70% cold ethanol before staining them with 1 µg/ml DAPI (Roche, Cat# 1023627001). To image fixed cells, we used an Olympus IX81 microscope equipped with a XM10 camera and a mercury lamp.

To apply osmotic and calcium stresses, we used a CellASIC ONIX2 system (EMD Millipore), controlled by a desktop computer through the software ONIX (EMD Millipore). To trap cells in the imaging chamber of a microfluidics plate (Y04C), we injected the exponentially growing cell culture to the chamber at a pressure of 34.5 KPa for 2 mins, followed by a continuous infusion of YE5s media at a pressure of 10KPa for another 30 mins. To apply hyper-osmotic stresses, we injected YE5s plus sorbitol into the chamber at a pressure of 10 KPa. Similarly, to apply hypo-osmotic stresses, we injected YE5s into the chamber at a pressure of 10 KPa after the chamber had been equilibrated with a continuous infusion of YE5s plus sorbitol for 30 mins.

We captured the kinetics of cytokinetic calcium spikes by taking time-lapse microscopy of the GCaMP-expressing cells at a frequency of 0.33 frame/sec without the Z-series. A complete spike is defined as one with a single peak rising from and decaying to the baseline level. For the constricting spikes, we examined the dividing cells with either two separating nuclei or a constricting ring.

### Image processing and analysis

We quantified the intracellular fluorescence of a GCaMP expressing cell as the total fluorescence intensity of the cell divided by its area. This value was used to represent the intracellular calcium level throughout the study. Cells were segmented manually, and their fluorescence intensities were quantified using average intensity projections of the Z-series. The intracellular fluorescence of the cells trapped in a microfluidics chamber was quantified by subtracting their combined fluorescence with the background fluorescence of the camera. Unless specified, we normalized the intracellular fluorescence of a cell against the baseline value, which is typically defined as the average of the five lowest values recorded during a time course. The microscopy images acquired during osmotic or calcium shocks were analyzed by quantifying two regions of interest in cytoplasm, each measuring 1 µm in diameter. The statistics tests were done using Excel. In most cases, two-tailed Student T-test was used. Standard deviations (S.D.) are used throughout the study to represent data deviations.

For quantification of the amplitudes and number of cytokinetic calcium spikes, we first segmented the dividing cells manually using NIH ImageJ. These videos are first pre-processed using a median filter to reduce noise. This non-linear filter helps smoothen the images while preserving high spatiotemporal resolutions. Following the noise removal, the background fluorescence signal from the cells is eliminated by subtracting the temporal average of each pixel in every frame. This process enables us to retain most of the flickering events, while eliminating the background fluorescence to a large extent. The average fluorescence intensity of a cell is then calculated by applying a binary mask on the images. Binarization of the median filtered images are performed by using Otsu’s thresholding method and applied to the background subtracted images. The average fluorescence intensity value at each time point is then recorded and used for analysis. Prior to identifying the peaks, we perform a temporal averaging of the signal, using two consecutive time points, to eliminate the small fluctuations within the data. Further thresholding of the signal is performed by nullifying the values lower than the temporal average of the entire signal. The temporal average is used as one of the criterions to distinguish the peaks from background fluctuations. This thresholding step removes the variation in the remnant background fluorescence signal inside the cells. The peaks or local maxima are detected by calculating the derivative at each point and identifying the points corresponding to the inversion of the derivatives from positive to negative. The location and average fluorescence intensity values of each individual peaks are then recorded. The video processing and analysis was performed using MATLAB (Mathworks, MA).

To measure the rate of the contractile ring closure, we determined the duration of the ring closure through analyzing the fluorescence kymograph of contractile ring. The closure rate was then calculated by dividing the initial circumference of the ring before the constriction by the duration. We used Image J (NIH) and custom-made macros for all the image analyses. The images were first corrected for X-Y drift using the Image J StackReg plug-in (Thevenaz et al., 1998) if necessary. Most plots were made with Kaleidagraph (Synergy, PA) except for the 3D plots which were made with OriginLab (OriginLab, MA). The figures were made with Canvas (ACDSee Systems).

## Acknowledgement

QC conceptualized the study, QC and AP designed and carried out the experiments, QC, AP, OS, and AR analyzed the data, QC and AP wrote the manuscript.

The authors would thank Debatrayee Sinha, Mamata Malla, Zachary Kreais and Somaiyeh Khoubafarin for their technical help. They would also like to thank our colleagues at the University of Toledo Song-Tao Liu, William Taylor and Rick Komuniecki for suggestions. They would thank Dimitris Vavylonis (Lehigh University) and Ann Miller (University of Michigan) for thoughtful discussions. This work has been supported by the University of Toledo startup fund (QC), NIH R15GM134496 (QC) and DeArce-Koch Memorial Fund (QC).

## Supplemental Materials

### Supplemental Figure legends

**Figure S1:**
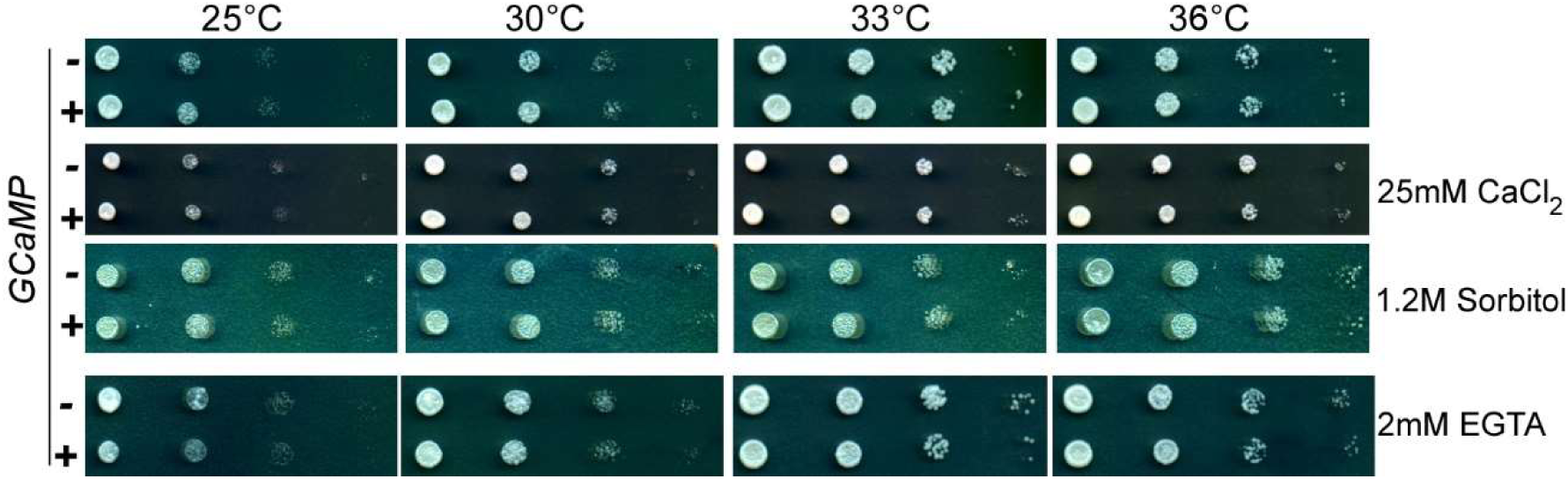
Expression of GCaMP does not affect fission yeast growth. Related to Fig. 1. Ten-fold dilution series of both GCaMP-expressing and the wild-type cells. The cells were inoculated at various temperatures on either YE5s (top), or YE5s plus 25mM CaCl_2_, or YE5s plus 1.2M sorbitol, or YE5s plus 2mM EGTA plates. GCaMP did not affect the yeast growth under any of the conditions. Similar results were obtained in two biological repeats.

**Figure S2:**
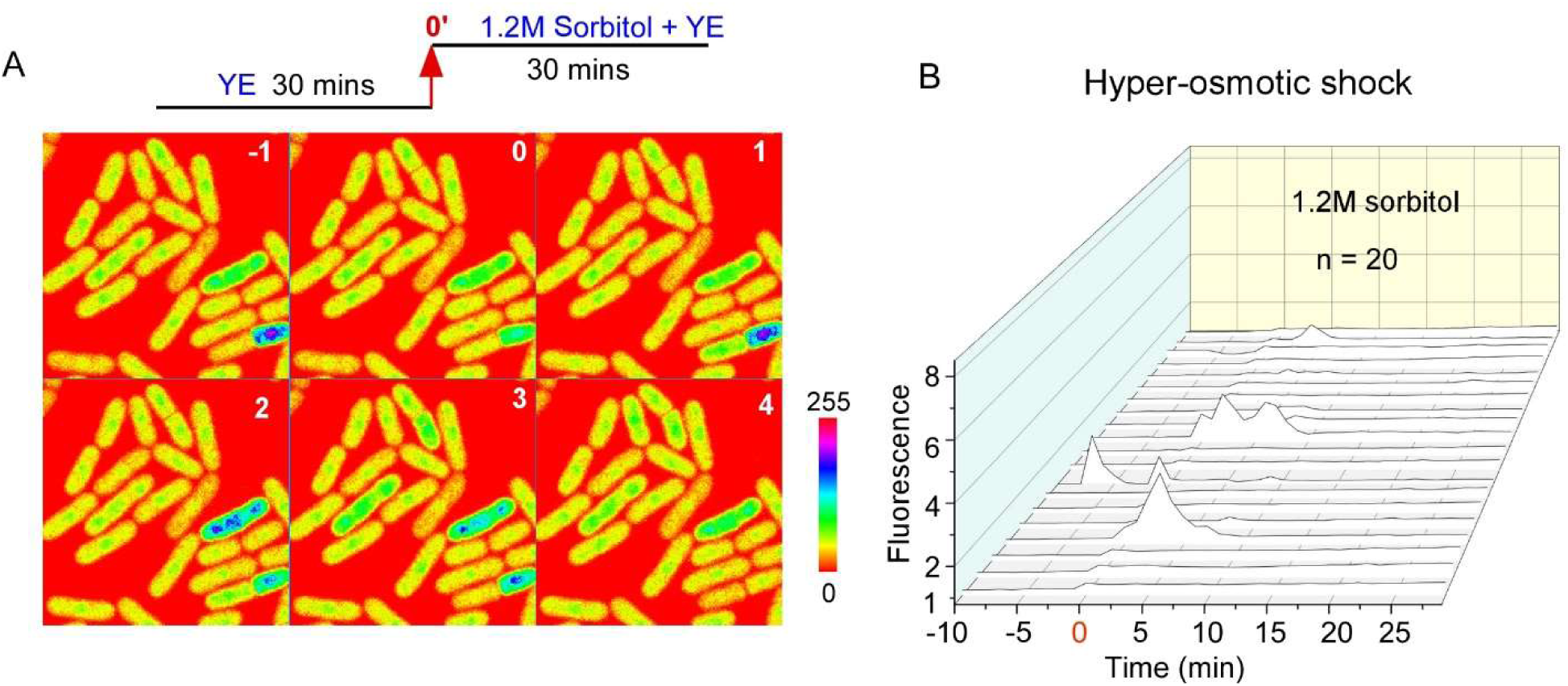
Hyper-osmotic shocks do not trigger calcium spikes. Related to Fig. 3. The cells were equilibrated with YE5s media for 30 mins in the microfluidic chamber before the infusion of YE5s plus 1.2M sorbitol (time zero). **(A)** Time-lapse micrographs (spectrum-colored) of the GCaMP expressing cells in a microfluidics chamber. Number: time in minutes after the shock. Unlike hypo-osmotic shock, hyper-osmotic shock did not trigger synchronized intracellular calcium spikes. Similar result was obtained in two biological repeats. **(B)** 3D line plots showing the time courses of the intracellular GCaMP fluorescence in twenty representative cells during the hyper-osmotic shock.

**Figure S3:**
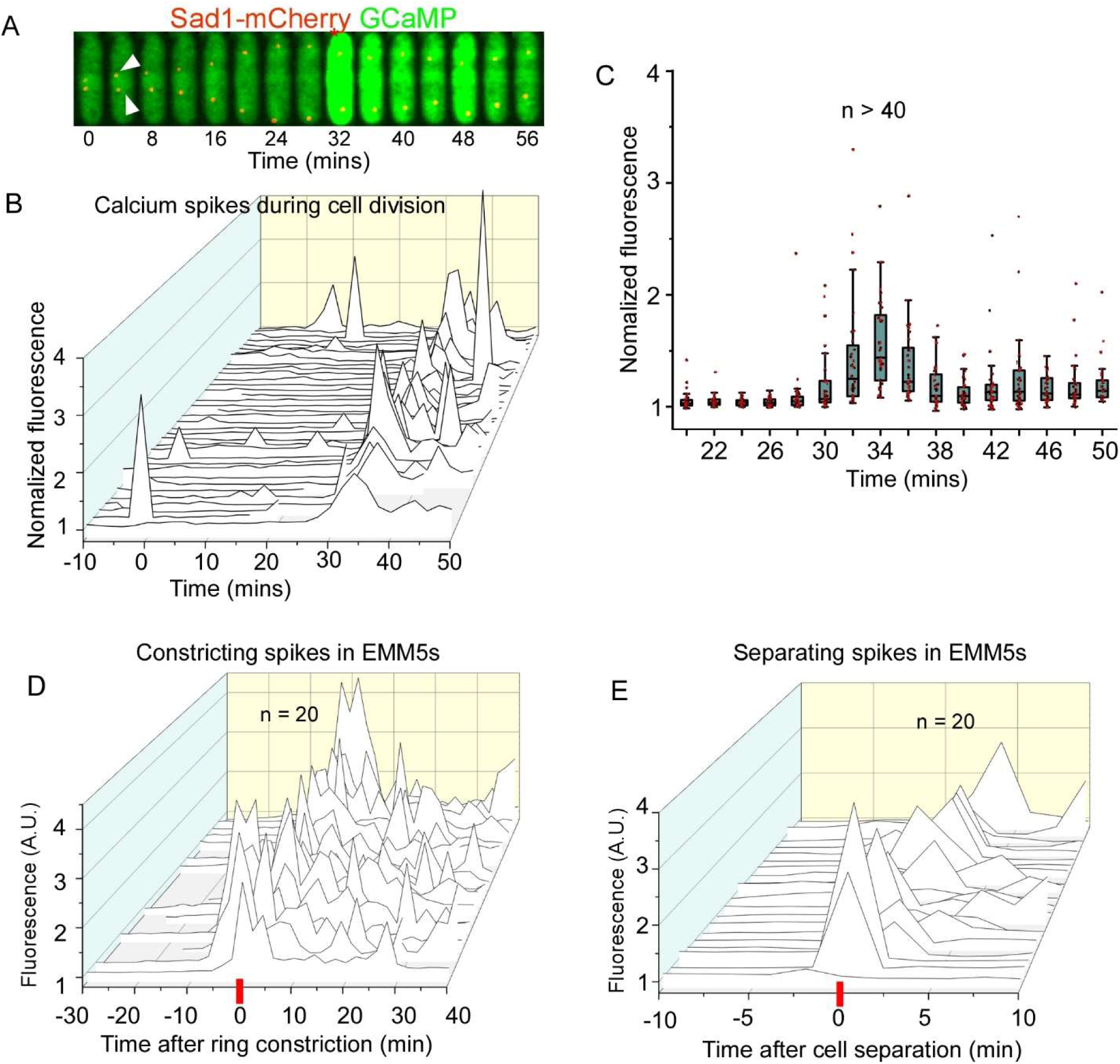
The calcium spikes in dividing cells. Related to Fig. 4, 5 and 6. **(A-C)** The intracellular calcium level increased during late mitosis. **(A)** Time-lapse fluorescence micrographs of a cell expressing both Sad1-mCherry (red) and GCaMP (green). Number: time in minutes after the SPB separation (arrowheads). Interval = 4 mins. The intracellular calcium level peaked at +32 mins (asterisk), at the start of telophase when two SPBs reached a maximum distance from each other. **(B)** 3D-line plots of the time courses of calcium level in mitotic cells (n = 36). The calcium level increased significantly starting ∼ 30 mins after the separation of SPBs. **(C)** Box plots of the calcium level of those cells shown in B. The average calcium level peaked at +34 min, after the separation of the SPBs. The data was pooled from two biological repeats. **(D-E)** The cytokinetic calcium spikes in EMM5s media. 3D-line plots of the calcium level during the ring constriction (D) and cell separation (E). Similar constricting and separating spikes are found in these cells, as those inoculated in YE5s.

**Figure S4:**
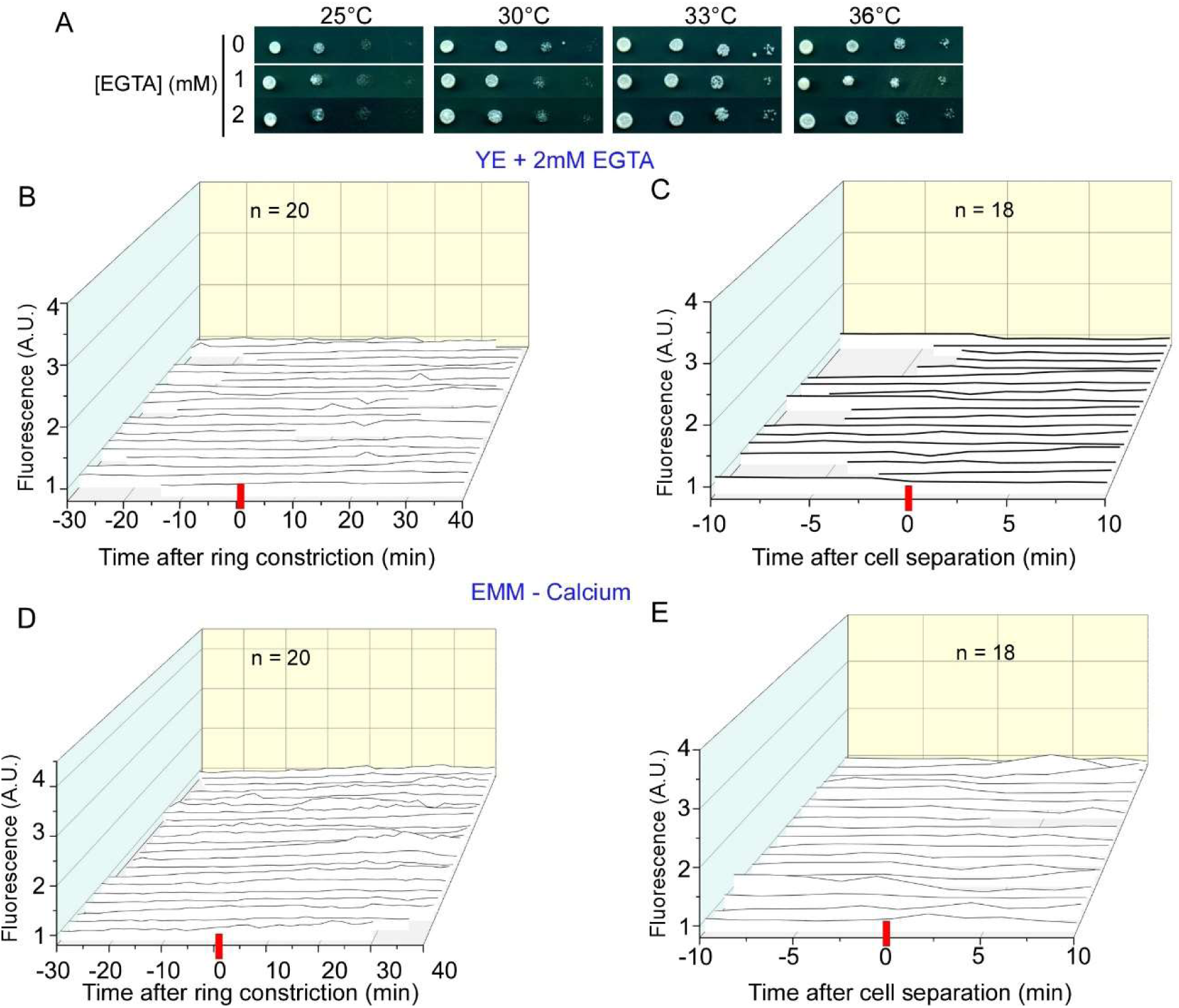
Depletion of extracellular calcium inhibits the cytokinetic calcium spikes. Related to Fig. 8. **(A)** Ten-fold dilution series of the wild-type yeast in YE5s plus 0-2 mM EGTA. Low concentration of EGTA didn’t inhibit cell growth. **(B-E)** 3D-line plots showing the time courses of the calcium level of the representative cells in either YE5s plus 2mM EGTA (B-C) or the calcium-free EMM media (D-E) during either the ring closure (B and D) or the cell separation (C and E). Depletion of extracellular calcium inhibited the cytokinetic calcium spikes. The data is pooled from two biological repeats.

**Figure S5:**
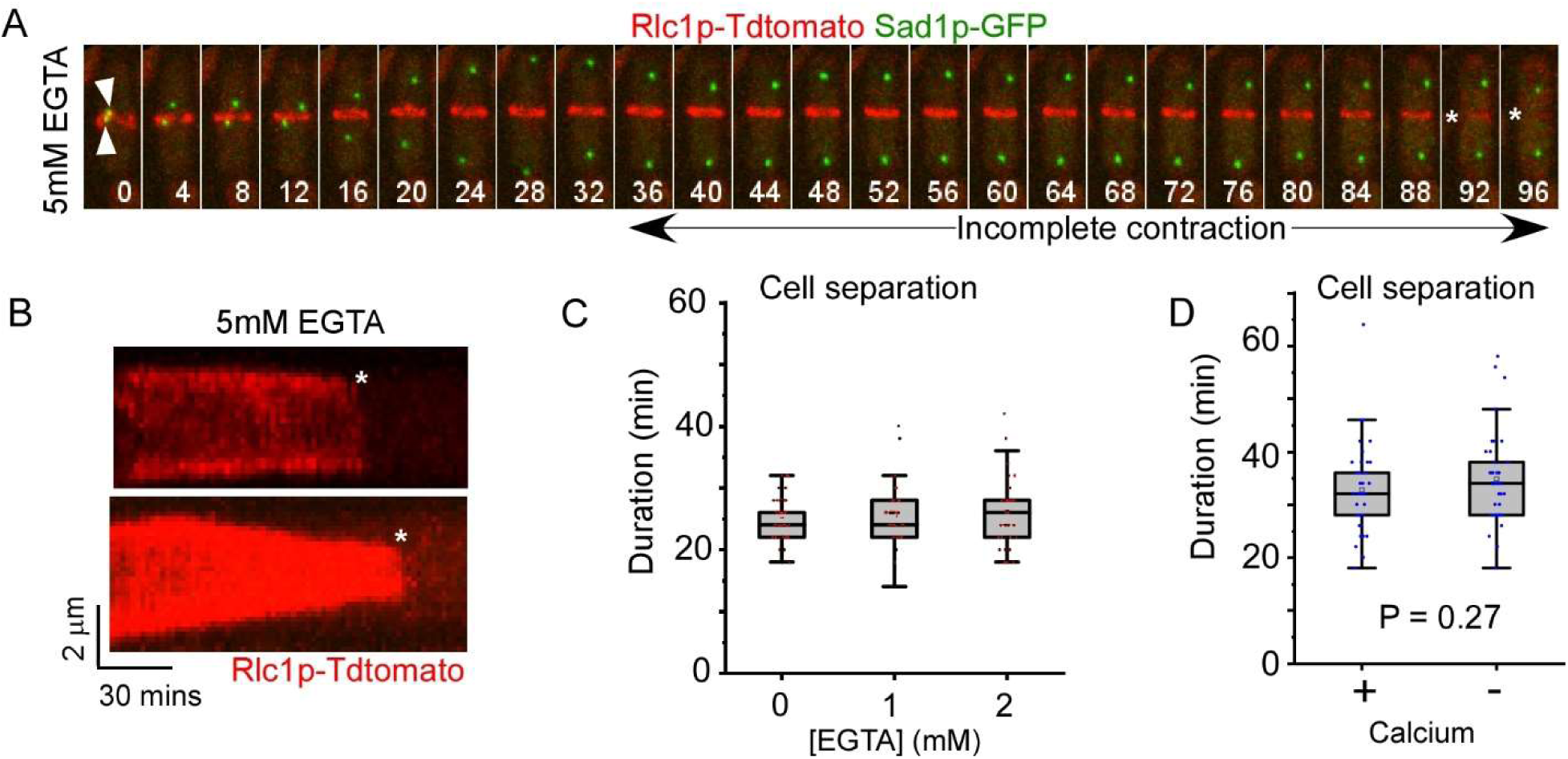
Depletion of extracellular calcium slowed down the contractile ring constriction. Related to Fig. 9. **(A-B)** Depletion of calcium by 5 mM EGTA reduces the integrity of the contractile ring. The cells expressed both Sad1p-GFP (green) and Rlc1p-tdTomato (red). **(A)** Time-lapse micrographs of a cell in YE5s supplemented with EGTA. The contractile ring failed to close completely before its disintegration (asterisk). Number: time in mins after the separation of SPBs (arrowhead). **(B)** Kymographs of two contractile rings that failed to constrict completely before their disintegration (asterisk). **(C-D)** Box plots showing the duration of cell separation when extracellular calcium was depleted. The separation was not significantly delayed.

### Supplemental Tables

**Table 1:** The list of yeast strains.

| Strains | Genotype | Source |
| --- | --- | --- |
| FY527 | *h- leu1-32 ura4-D18 his3-D1 ade6-M216* | Lab stock |
| FY528 | *h+ leu1-32 ura4-D18 his3-D1 ade6-M210* | Lab stock |
| QC-Y941 | *h- leu1::KanMX6-Padh1-GCaMP6s* | This study |
| QC-Y989 | *h- sad1-mCherry-NatMX6 leu1::Padh1-GCaMP6s-mCherry* | This study |
| QC-Y1004 | *h+ rlc1-tdTomato-NatMX6 leu1::Padh1-GCaMP6s-mCherry* | This study |
| QC-Y1068 | *h- leu1::Padh1-GCaMP6s-mCherry-ura4+* | This study |
| QC-Y274 | *h- rlc1-tdTomato-NatMX6 sad1-mGFP-KanMX6* | Lab stock |
| QC-Y1149 | *h? pmr1::kanMX6 rlc1-tdTomato-natMX6 sad1-mEGFP-KanMX6* | This study |

**Table 2:**
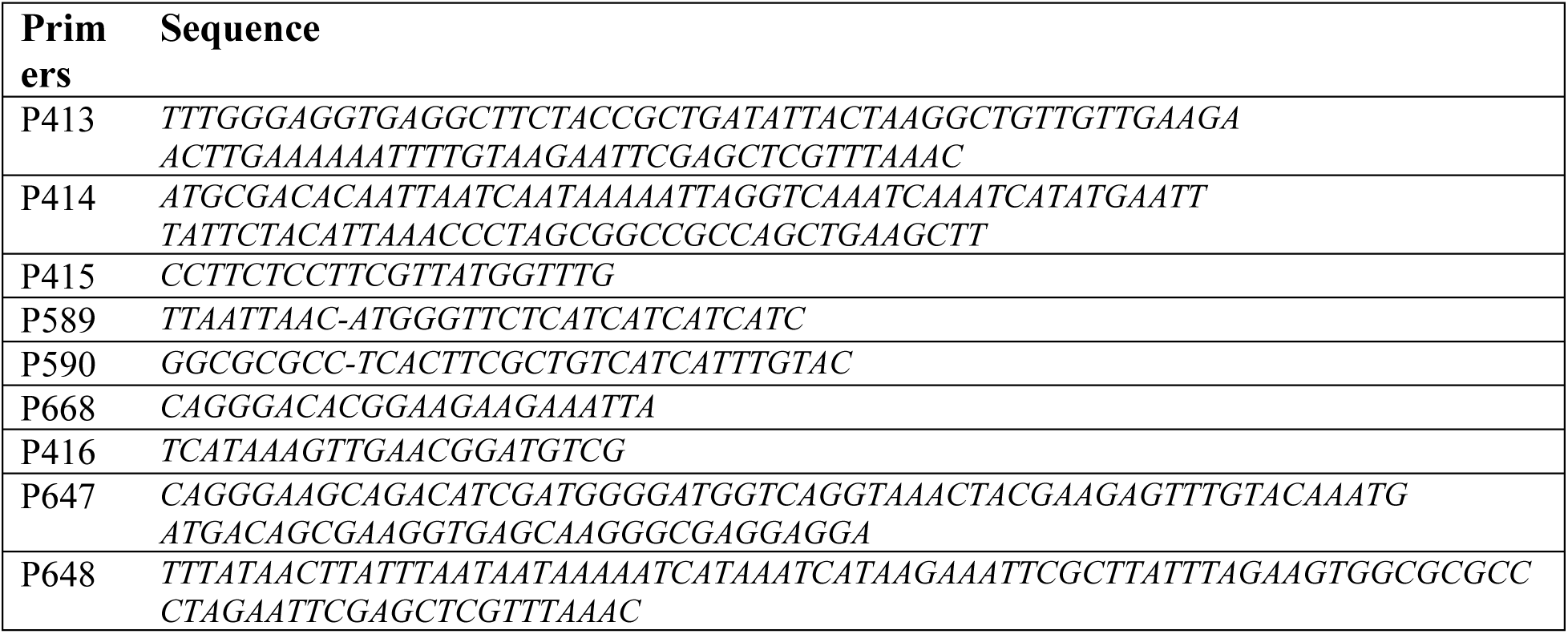
The list of DNA primers.

### Supplemental movies

**Movie S1: The calcium homeostasis of fission yeast reported by GCaMP-based imaging (related to Fig. 2)**. Time-lapse movie (60 mins) of GCaMP expressing cells in YE5s. Number: time in minutes.

**Movie S2: The calcium spikes stimulated by hypo-osmotic shock (related to Fig. 3)**. Time-lapse movie of the GCaMP expressing cells in a microfluidic chamber. At time zero, the infusing media was switched from YE5s plus 1.2M sorbitol to just YE5s. This hypo-osmotic shock (time zero) triggered synchronized calcium transients. Number: time in minutes.

**Movie S3: The constricting calcium spike during cytokinesis (related to Fig. 4)**. Time-lapse movie of a dividing cell expressing both GCaMP (green) and Rlc1p-tdTomato (red). Number: time in minutes. The calcium spike initiated as the ring constriction started (time zero) and peaked at +2 min. Number: time in minutes.

**Movie S4: The separating calcium spike during cytokinesis (related to Fig. 5)**. Time-lapse movie of a GCaMP expressing cell during the cell separation. The calcium spike followed the separation (time zero), oscillating between the two daughter cells. Number: time in minutes.

**Movie S5: Cytokinesis in the calcium depleted media (related to Fig. 9)**. Time-lapse movie of two cells in either YE5s (left) or YE5s plus 2mM EGTA (right). The cells expressed both GCaMP (green) and Rlc1p-tdTomato (red). EGTA inhibited the cytokinetic calcium spikes. The contractile ring constricted more slowly in the presence of EGTA. Number represents time in minutes after the start of the ring constriction (time zero).

